# The hippocampus shows an own-age bias during unfamiliar face viewing

**DOI:** 10.1101/2021.02.15.431110

**Authors:** Joshua D. Koen, Nedra Hauck, Michael D. Rugg

**Affiliations:** Department of Psychology, University of Notre Dame; Center for Vital Longevity and School of Behavioral and Brain Sciences, University of Texas at Dallas; School of Psychology, University of East Anglia, UK

## Abstract

The present study investigated the neural correlates of the own-age bias for face recognition in a repetition suppression paradigm. Healthy young and older adults viewed upright and inverted unfamiliar faces. Some of the upright faces were repeated following one of two delays (lag 0 or lag 11). Repetition suppression effects were observed in bilateral fusiform cortex. However, there were no significant effects indicating an own-age bias in repetition suppression. The absence of these effects is arguably inconsistent with perceptual expertise accounts of own-age biases in face processing. By contrast, the right anterior hippocampus showed an own-age bias (greater activity for own-than other-age faces) when viewing an unfamiliar face for the first time. Given the importance of the hippocampus for episodic memory encoding, we conjecture that the increased hippocampal activity for own-age relative to other-age faces reflects differential engagement of neural processes supporting the episodic encoding of faces and might provide insight into the neural underpinnings of own-age biases in face recognition memory.

## Introduction

The ways in which we evaluate, attend to, and remember human faces are susceptible to own-age biases (e.g., Bartlett and Fulton, 1991; Ebner, 2008; Wiese et al., 2008; Rhodes and Anastasi, 2012). Notably, both young and older adults show better recognition memory for own-age faces (Rhodes and Anastasi, 2012). These findings have motivated research that has demonstrated own-age biases in neural correlates of face processing (e.g., Ebner et al., 2011a, 2011b, 2013; Neumann et al., 2015; Wiese et al., 2008; Wolff et al., 2012; Ziaei et al., 2019; for review, see Wiese et al., 2013). A growing body of fMRI research aimed at identifying the neuroanatomical correlates of the own-age bias has reported BOLD signal increases for own-age versus other-age faces in the amygdala, medial prefrontal cortex, orbitofrontal cortex, and insula across a variety of conditions. (Ebner et al., 2013, 2011a; Wright et al., 2008; Ziaei et al., 2019; see also Golarai et al., 2017). Although the above studies did not report own-age effects in the fusiform gyrus, a canonical face-processing region, Golarai et al. (2017) recently reported such effects in the fusiform gyrus in both children (7-10 years) and young adults (18-40 years).

The present study had two goals. First, building on prior research, we investigated if own-age biases are present in repetition suppression effects elicited by unfamiliar faces. Repeating a face elicits a ‘repetition suppression’ effect whereby the BOLD response is reduced for repeated compared to first presentation faces (see Henson, 2016; Henson and Rugg, 2003). Repetition suppression in regions such as the occipital and fusiform face areas are proposed to reflect modulation of processes contributing to the identification of individual faces (Goh et al., 2010; Hermann et al., 2017). This proposal aligns with perceptual expertise theories of the own-age bias arguing that own-age faces are processed more efficiently and are better individuated than other-age faces due to more extensive experience with own-age peers. Second, we aimed to conceptually replicate prior studies (Wright et al., 2008) by examining own-age biases when viewing unfamiliar faces without employing a task that may induce strategy differences between young and older adults.

## Materials and Methods

### Ethics Statement

This study was approved by Institutional Review Board of the University of Texas at Dallas and University of Texas Southwestern Medical Center. All participants provided written informed consent prior to participation.

### Participants

A sample of 24 young and 26 older participants contributed to the analyses reported here. Participants were recruited from the University of Texas at Dallas and the greater Dallas metropolitan area and were financially compensation for their time ($30/hour). The sample sizes were determined by the requirements of a primary experiment in which the aim was to obtain usable data from 24 young and 24 older adults for a memory encoding task (Koen et al., 2019). These were the largest samples that could be accommodated given the resources available for the project. The higher number of participants contributing to the analyses reported here results from the fact some participants were excluded from the primary experiment but retained in the present study.

All participants were right-handed, reported having normal or corrected-to-normal vision and had no contraindications to MRI scanning. Exclusion criteria included a history of cardiovascular disease (other than treated hypertension), diabetes, psychiatric disorder, illness, or trauma affecting the central nervous system, substance abuse, and self-reported current or recent use of psychotropic medication or sleeping aids. All participants were considered cognitively normal as determined by performance on a test battery (Table 1; for details of the battery see Koen et al., 2019). All participants scored 27 or more on the Mini-Mental State Examination (MMSE; Folstein et al., 1975) and no more than 1.5 standard deviations below age-normalized scores on any one memory measure or on two (or more) non-memory measures. Data from an additional young adult male and one older adult male were excluded due to excessive in-scanner motion (> 8 mm maximum frame-wise displacement) during the task.

**Table 1.** Demographic and neuropsychological test data for young and older adults.

|  | Young Adults | Older Adults | *p*-value |
| --- | --- | --- | --- |
| N | 24 | 26 |  |
| Age | 23.04 (3.46) | 69.19 (3.27) |  |
| Sex (M/F) | 12/12 | 14/12 |  |
| Education | 15.92 (2.22) | 16.96 (2.24) | .104 |
| MMSE | 29.54 (0.59) | 29.35 (0.94) | .378 |
| CVLT Short Delay – Free | 13.08 (1.79) | 10.38 (3.16) | < .001 |
| CVLT Short Delay – Cued | 13.67 (1.81) | 11.67 (3.57) | .016 |
| CVLT Long Delay – Free | 13.54 (2.06) | 10.23 (3.27) | < .001 |
| CVLT Long Delay – Cued | 14.12 (1.62) | 11.92 (2.78) | .001 |
| CVLT Recognition – Hits | 15.42 (0.83) | 15.00 (0.98) | .110 |
| CVLT Recognition – False Alarms | 0.46 (0.66) | 3.08 (2.53) | < .001 |
| Logical Memory I | 30.62 (4.95) | 26.00 (5.50) | .003 |
| Logical Memory II | 28.12 (5.78) | 22.62 (6.03) | .002 |
| Digit Span Total<sup>1</sup> | 21.04 (4.53) | 17.77 (2.55) | .004 |
| SDMT | 65.38 (13.99) | 46.42 (7.74) | < .001 |
| Trails A (secs) | 21.43 (7.97) | 31.55 (10.76) | < .001 |
| Trails B (secs) | 47.54 (19.53) | 70.52 (24.87) | .002 |
| F-A-S Total | 48.29 (10.97) | 45.50 (11.42) | .382 |
| Category Fluency (Animals) | 24.58 (5.67) | 21.00 (4.66) | .020 |
| WTAR (Raw) | 41.42 (3.44) | 43.08 (4.78) | .163 |
| Raven's (List 1) | 11.08 (.97) | 9.19 (2.50) | .001 |
| Visual Acuity (logMar)<sup>2</sup> | -.11 (.10) | .06 (.11) | < .001 |

*Note.* Standard deviations are reported in parentheses. The *p*-values were obtained from unequal-variance independent *t*-tests comparing young and older adults. <sup>1</sup>Digit span total equals the sum of forward and backward span. <sup>2</sup>Lower logMAR scores indicate better visual acuity. MMSE = Mini-mental State Exam; CVLT = California Verbal Learning Test II; SDMT = Symbol-Digit Modalities Test; WTAR = Wechsler Test of Adult Reading

### Materials and Procedure

The critical stimuli comprised 208 faces from the CAL/PAL database (Ebner, 2008; Minear and Park, 2004). All face stimuli were 640×480 pixels with a grey background. Half of the face stimuli depicted younger adults (age range: 18-34 years) and half depicted older adults (age range: 60-91 years). The 104 stimuli of each face age group were randomly assigned to the lag 0 repeat (24 faces), lag 11 repeat (24 faces), control (32 faces) and inverted (24 faces) conditions, i’hc stimuli were further split into two lists, with half of the stimuli from each of the face age by trial type conditions. There were an equal number of male and females faces in each condition. An additional 8 images were used in a practice task completed outside of the scanner. Stimuli were presented to participants via a mirror mounted to the head coil. Cogent software (www.vislab.ucl.ac.uk/cogent_2000.php) as implemented in Matlab 2011b (www.mathworks.com) was used for stimulus control and response logging.

The experimental procedure is depicted in Figure 1. Participants were shown a series of faces (1 sec duration followed by a 1.25 sec white fixation cross) and were instructed to press a key with their right index finger whenever an inverted face was presented. Speed was emphasized on the button press to inverted faces. There were 48 null trials dispersed throughout the block to jitter the stimuli, with no more than one null trial occurring consecutively. Responses were recorded until the beginning of the next trial.

**Figure 1.**
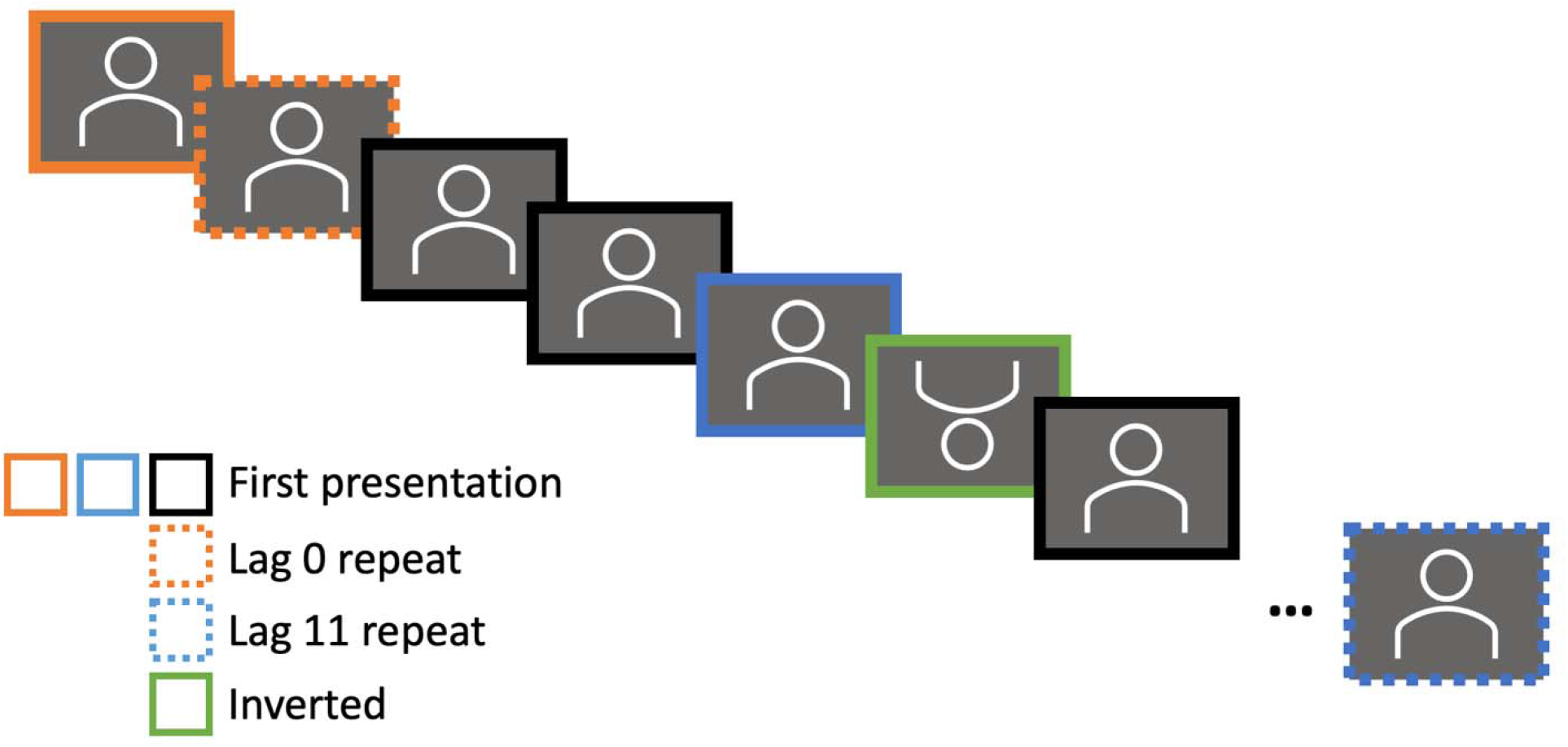
Schematic of the face repetition task. Participants saw a series of young and older adult faces in a face inversion task. Participants were instructed to make a button press if the face was inverted, and otherwise to simply view the faces. Each face was shown for 1 second and followed by an inter-trial interval (fixation cross) for 1.25 seconds. There was a total of 8 conditions formed by crossing Face Age (young, older) and Trial Type (first, lag 0 repeat, lag 11 repeat, inverted). Additional details are reported in the Materials and Methods section.

Each session of the task comprised 48 null trials and 152 face trials (80 first presentation trials, 24 lag 0 repeats; 24 lag 11 repeats, 24 inverted faces; equally split across faces depicting young and older adults). The decision to employ only two lags was driven by time constraints, the rather limited number of stimuli, and our attempt to maximize the number of trials in ‘short’ and ‘long’ repetition lags. Inverted faces were presented once only, and no more than two inverted trials occurred in succession. Faces assigned to the repeat condition were presented twice with the repetition occurring either on the immediately succeeding trial (lag 0) or after 11 intervening trials (lag 11). Faces in the control condition were presented on only one occasion. Control faces and the first presentations of repeated faces were collapsed into a single ‘first’ face condition.

### Behavioral Data Analysis

The log-transformed reaction times to inverted face stimuli receiving a correct response were submitted to a linear mixed model using the *mixed()* function from the *afex* package (Singmann et al., 2021) in R (R Core Team, 2021). The model included fixed effect terms age group (young versus older), face age congruency (own age versus other age), and their interaction. Same age faces were those faces from the same age group as the participant (e.g., young faces were coded as same age faces for young adults) whereas different age faces were those face stimuli depicting individuals not in the participants age group (e.g., for young adults, older faces were coded as different age faces). The model included random effects of participant, specifically a random intercept and random slope of face age congruency. Degrees of freedom were estimated using the Satterthwaite (1946) approximation. Bayes factors for the alterative (BF_10_) hypothesis were computed using the *ImBF*function from the *Bayes* Factor package(Morey and Rouder, 2018) and *bayesfactor_inclusion* function from the *bayestestR* package (Makowski et al., 2019).

### MRI Data Acquisition

MRI data were acquired with a 3T Philips Achieva MRI scanner (Philips Medical Systems, Andover, MA, USA) equipped with a 32-channel receiver head coil. Functional images were acquired with a blood oxygenation level dependent (BOLD), T2*-weighted echoplanar imaging (EPI) sequence (SENSE factor = 1.5, flip angle = 70°, 80 × 80 matrix, FOV = 240 mm x 240 mm, TR = 2000 ms, TE = 30 ms, 34 ascending slices, slice thickness = 3 mm, slice gap = 1 mm), and were oriented parallel to the AC-PC line. Five “dummy” scans were acquired at the start of each fMRI session and discarded to allow for equilibration of tissue magnetization. A total of 264 functional volumes were acquired during each of the two task runs, for a total of 528 brain volumes. T1-weighted images (MPRAGE sequence, 240 × 240 matrix, 1 mm isotropic voxels) were acquired for anatomical reference. Note that this task was completed prior to the encoding phase of a memory study that comprised object and scene images (Koen et al., 2019).

### fMRI Data Preprocessing and Analysis

The functional data were preprocessed with Statistical Parametric Mapping (SPM12, Wellcome Department of Cognitive Neurology, London, UK) implemented in Matlab 2017b (The Mathworks, Inc., USA). The images were reoriented, subjected to a two-pass realignment procedure, whereby images were initially realigned to the first image of a session and then realigned to the mean EPI image, and then corrected for slice acquisition time differences using sinc interpolation with reference to the middle slice. Finally, images were spatially normalized to a study specific EPI template (de Chastelaine et al., 2011) and smoothed with an 8mm full-width at half-maximum kernel.

The fMRI data were analyzed with a two-stage random effects model. First level GLMs modeled neural activity as a delta function convolved with a canonical HRF for the 8 event types formed by crossing face age and the four item types (first, lag0 repeat, lag11 repeat, inverted face). The first level GLM also included the 6 realignment parameters and session specific means as nuisance variables.

The second-level group analyses were performed by submitting the 8 first-level beta maps to a 2 (age group) by 8 (condition) mixed ANOVA using the factorial ANOVA module in SPM12. The 8 levels of the condition factor were those described above for the first-level models, with one exception. Like the analysis of the reaction time data, face age was coded as a face age congruency factor with own-age and other-age faces. Planned contrasts were conducted to identify voxels showing significant effects of repetition, face age congruency, and to examine whether these effects interacted with participant age or lag. Additionally, effects of face inversion were examined. Effects were deemed significant if they survived p < .001 with a cluster-wise correction *(p* < .05, FWE) based on Gaussian Random Field theory or if the peak voxel of a cluster survived *p* < .05 FWE voxel-level correction. The rationale for using both approaches was to avoid Type-II errors both for large clusters with (relatively) weak mean activation and for more focal clusters with a strong peak response.

As noted in the Results, we further probed the whole-brain findings by extracting the mean beta response from 5mm spheres centered on the peak effects. These values were subjected to mixed factor ANOVAs using the JASP software (JASP Team, 2020). Bayes factors for effects favoring the alternative (BF_10_) hypothesis are reported for these follow-up analyses.

## Results

### Behavioral Data

Participants were accurate in identifying inverted faces (≥ 98%) and very rarely false alarmed to upright faces (≤ .3%). The analysis of reaction times revealed null effects of age group, *F*(1,47.98) = 1.93, *p* = .171, partial-η^2^ = .04, BF_10_ = 0.693, face age congruency, *F*(1, 47.87) = 0.003, *p* = .958, partial-η^2^ = .00, BF_10_ = 0.069, and the interaction between the two variables, *F*(1, 47.87) = 1.36, *p*= .250, partial-η^2^ = .03, BF_10_ = 0.134.

### Face Inversion Effects

We first examined the effects of face inversion by contrasting the activity elicited by upright (first presentation only) and inverted faces. There was an increase in the BOLD signal for inverted relative to upright races across a wide swath of the cortex with peaks primarily in the frontal and temporal cortices, as well as the cerebellum. No voxels survived our statistical thresholds when looking for voxels showing elevated BOLD signal for upright compared to inverted faces. An additional contrast examining age differences in the face inversion effect identified clusters in the right pre- and postcentral gyri showing smaller face inversion effects for older relative to younger adults. No significant clusters demonstrated an interaction between face inversion and face age congruency (own vs. other age faces).

### Repetition Suppression

The first planned contrast identified voxels showing effects of face repetition. No clusters demonstrated significant repetition enhancement effects. However, clusters showing significant repetition suppression effects were identified in bilateral fusiform gyrus (Table 2 and Figure 2A-B). The suppression effects did not differ significantly between young and older adults, between repetition lags, or between own-age and other-age faces according to follow-up analyses that exclusively masked the repetition suppression contrast with interaction terms involving repetition and the other factors (exclusive mask threshold set at *p* < .10, uncorrected). The absence of age group differences in repetition suppression effects in the fusiform gyrus is consistent with some prior fMRI findings (Goh et al., 2010). However, the lack of an effect of repetition lag is inconsistent with previous research on face repetition (Henson et al., 2000; for related findings, see Nagy and Rugg, 1989); this inconsistency might be due to the relatively short lag in our ‘long-lag’ condition (maximum of 11 intervening trials before a face repeat) compared to prior studies.

**Figure 2.**
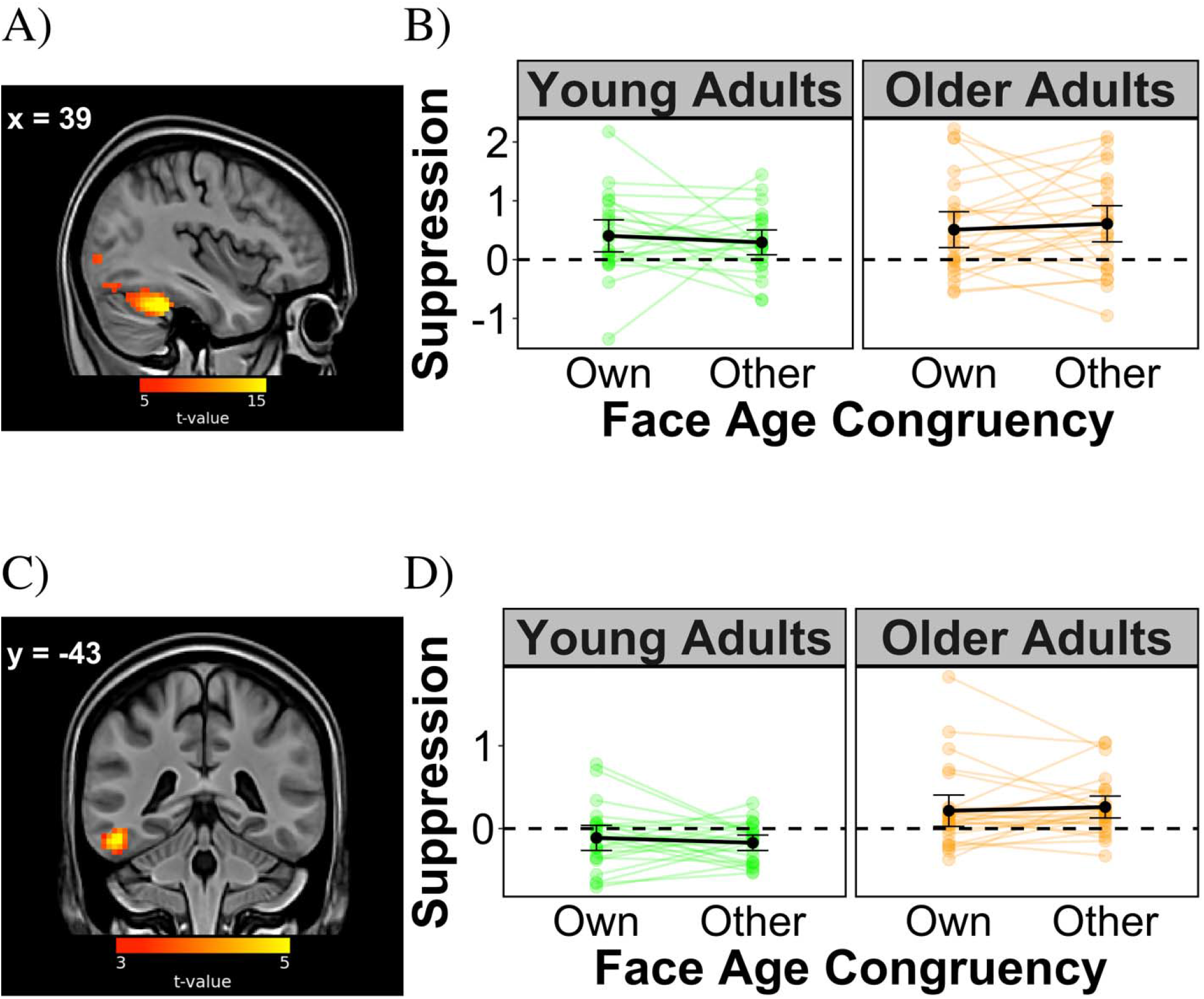
Age-group invariant (top panels) and age-group dependent (bottom panels) face repetition suppression effects. (A) A cluster in the right fusiform gyrus (x = 39, y = −43, z = −28) showing suppression of the BOLD signal for repeated faces compared to the first presentation of faces in both young and older adults. A similar pattern was observed in the left fusiform gyrus (not shown, see Table 1). (B) Repetition suppression estimates (first minus repeated presentation) extracted from a 5mm sphere centered on the right fusiform gyrus peak voxel. (C) A cluster in the left inferior temporal cortex (x = 39, y = −43, z = −28) showing larger repetition suppression effects for older relative to younger adults. (D) Repetition suppression estimates extracted from a 5mm sphere centered on the left inferior temporal cortex peak voxel. The interaction in this region is driven by the combination of repetition suppression in older adults and repetition enhancement in younger adults. (A) and (C) are shown at p < .001, uncorrected, for visualization purposes, and depicted in neurological orientation (right is right). In (B) and (D), the solid black circles represent condition means, and the green and orange points depict data from individual participants. Error bars reflect the 95% confidence intervals computed from the standard error of the observed data with custom code.

**Table 2.**
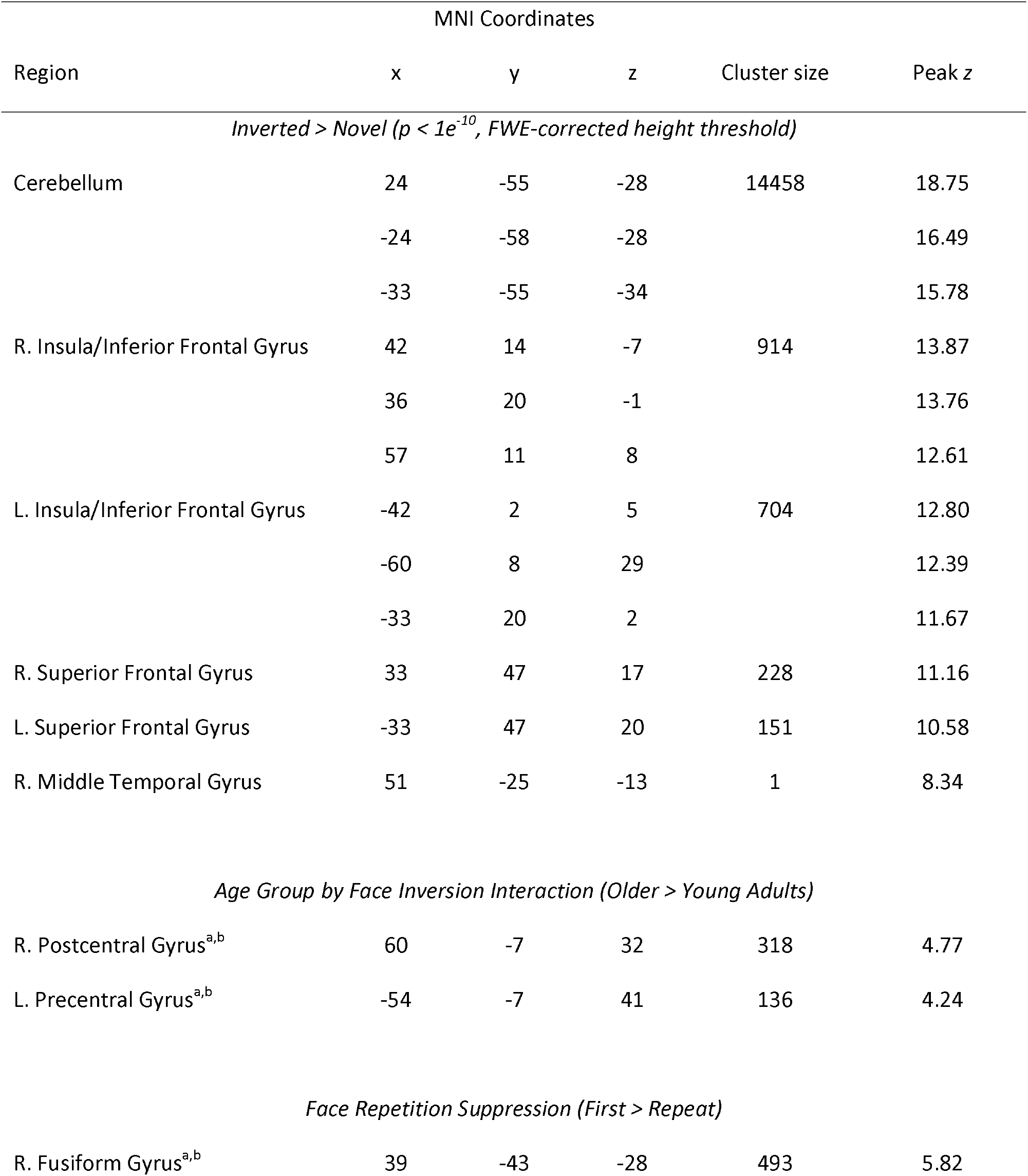
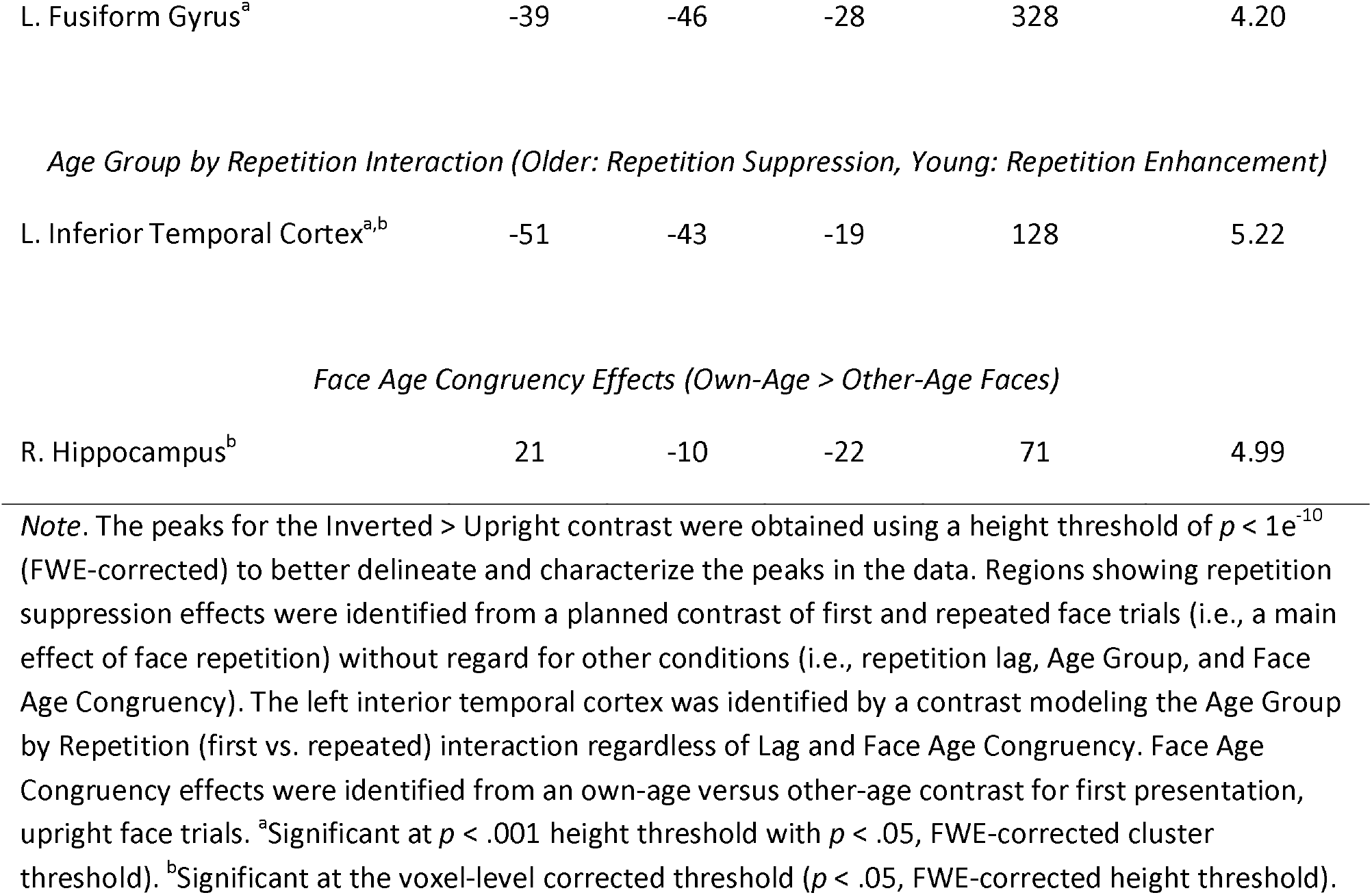
Regions showing effects of face repetition and age congruency.

We next conducted planned contrasts aimed at identifying voxels where face repetition effects differed in magnitude between young and older adults (i.e., regions showing an Age Group by Repetition Interaction, collapsed across lag). This contrast identified a cluster in the left interior temporal gyrus that showed repetition suppression effects in older but not young adults (Figure 2C-D). Repetition suppression indices (i.e., first minus repeat) extracted from a 5mm sphere centered on the peak voxel manifesting the interaction revealed a significant repetition suppression effect for repeated faces in older adults, *t*(25) = 3.24, *p* = .003, Cohen’s *d* = 0.635, BF_10_ = 11.93, but repetition *enhancement* for repeated faces in the younger adults, *t*(23) = 2.85, *p* = .009, Cohen’s *d* = 0.582, BF_10_ = 5.27.

A further planned contrast found no evidence that face repetition effects interacted with repetition lag or face-age congruency.

### Own-Age biases during novel face viewing

We next conducted a planned contrast between the first presentation trials of upright, own-age and other-age faces to identify voxels showing an own-age bias. Repeated faces were excluded to mitigate the potentially confounding effects of familiarity or other repetition-related processes on own-age bias effects, allowing us essentially to mirror the contrast employed to identify own-race effects reported by Brown et al. (2017). The above-described contrast identified a cluster of voxels demonstrating a face age congruency effect in the right anterior hippocampus (Table 2 and Figure 3). The cluster demonstrated elevated BOLD activity for own-age relative to other-age faces. This effect did not significantly differ in magnitude between young and older adults based on the outcome of an exclusive mask (at *p* < .10) of the above contrast with the interaction contrast between age group and face age congruency. Converging with this finding, a conjunction analysis performed by inclusively masking the own-age versus other-age contrasts conducted separately for young and older adults (each thresholded at *p* < .01) revealed a cluster that overlapped with that identified in the initial analyses.

**Figure 3.**
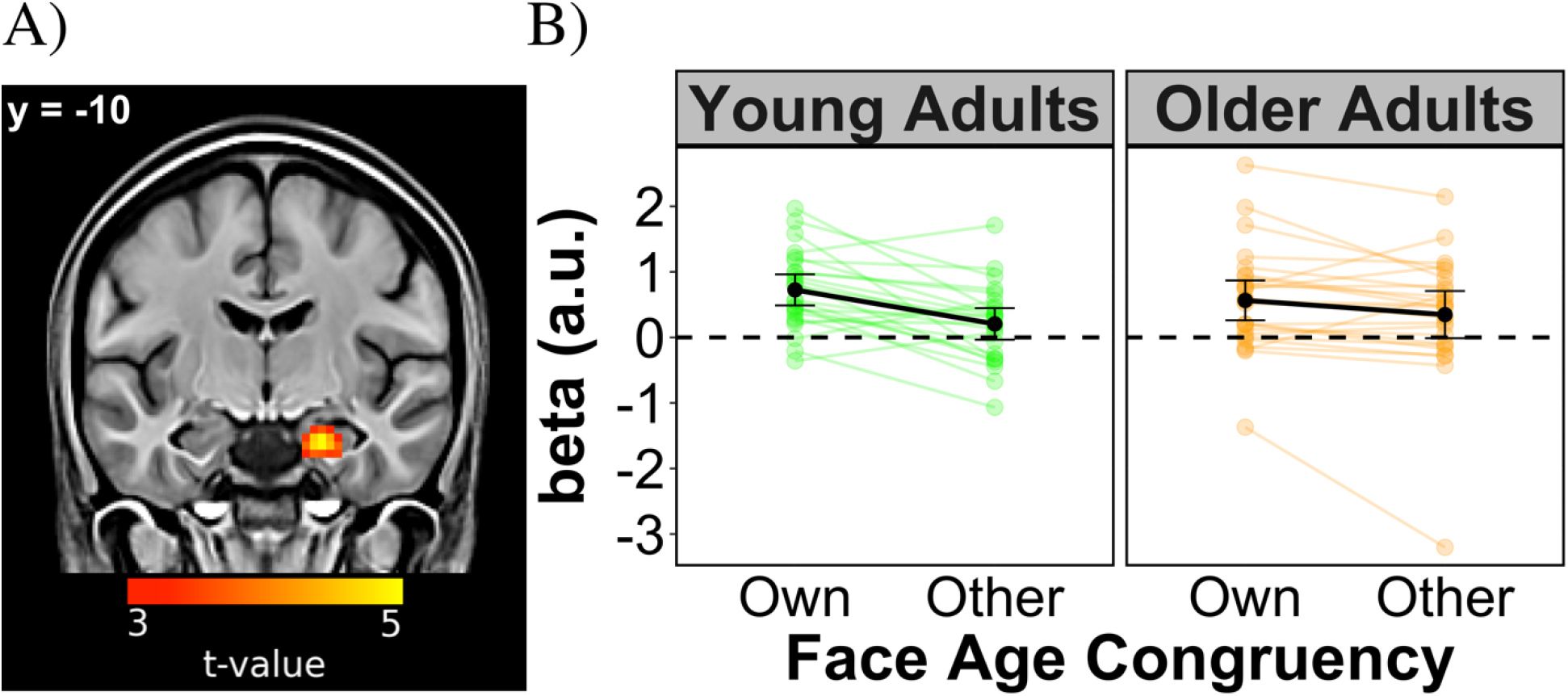
The BOLD signal in the right hippocampus was greater when young and older adults viewed faces belonging to their own age group relative to other age faces. (A) Viewing the first repetition of novel faces was associated with an own-age bias in the right anterior hippocampus (x = 22, y = −10, z = −24). The image is shown at a threshold of at p < .001 uncorrected, for visualization purposes. (B) The beta values extracted from a 5mm sphere centered on the peak coordinate in the right hippocampus. The solid black circles represent condition means, and the green and orange points depict data from individual participants. Error bars reflect the 95% confidence intervals computed from the standard error of the observed data with custom code.

Follow-up analyses investigated whether face repetition moderated the own-age bias effect observed in the right anterior hippocampus. We conducted a 2 (age group) by 3 (face condition) by 2 (face age congruency) mixed ANOVA on beta estimates for the three upright face trial types (first, lag 0, and lag 11) from a 5mm sphere centered on the peak of right anterior hippocampus cluster (x = 22, y = − 10, z = −24). Note that because the main effect of face age congruency is not independent of the contrast used to define the original face age effect, the effect is circular and therefore not considered here. The ANOVA revealed null effects for the two-way interaction between face age congruency and age group, *F*(1, 47) = 1.55, *p* = .219, partial-η^2^ = .143, BF_10_ = 0.308, for face condition, *F*(1.85, 86.85) = 1.21, *p* = .288, partial-η^2^ = .025, BF_10_ = 0.147, and for the three-way interaction, *F*(1.98, 92.93) = 0.645, *p* = .527, partial-η^2^ = .143, BF_10_ = 0.176. It is noteworthy that the Bayes factors provided moderate to strong evidence favoring null effects (BF_10_ < .333) according to the definitions proposed by Jeffreys (Jeffreys, 1939; Kass and Raftery, 1995). In short, these null findings suggest that none of factors included in the ANOVA significantly moderated the own-age bias effect observed in the right anterior hippocampus.

## Discussion

The present study examined whether face age moderated repetition suppression effects elicited by unfamiliar faces and whether there were any own-age biases in neural activity when viewing first presentations of unfamiliar faces. There were three main findings. First, although we identified significant age-invariant repetition suppression effects in bilateral fusiform gyrus, we did not identify any clusters where suppression effects demonstrated an own-age bias. These null findings are arguably inconsistent with perceptual expertise accounts of the own-age bias (Tanaka and Pierce, 2009; Valentine, 1991; Wiese et al., 2013). Such accounts propose that own-age faces are more efficiently processed than other-age faces due to more extensive experience with own-age peers. If face repetition suppression effects reflect modulation of processes contributing to the identification of individual faces (Hermann et al., 2017), then we might expect that own-age faces would elicit a greater suppression effect than other-age faces, which we did not observe. However, one limitation of the present design is that we only examined repetition effects for identical repeats and did not include a condition that could examine ‘release from suppression’ (e.g., Goh et al., 2010; Reggev et al., 2020). Recently, Reggev et al. (2020) argued in favor of a perceptual expertise account of own-race biases based on findings of greater release from suppression for own-race relative to other-race faces. Future research employing designs that allow measurement of release from suppression is needed to pursue this issue.

Second, we identified a cluster in left inferior temporal cortex that demonstrated repetition suppression effects for older adults but repetition enhancement effects in young adults. This region has been previously reported to demonstrate repetition suppression effects for familiar (e.g., yellow banana) but not novel concepts (e.g., purple banana) (Reggev et al., 2016). Given that familiar concepts are accrued through experience, it is possible that the present interaction reflects age group differences in exposure to faces (Golarai et al., 2017; for a related discussion, see Koen and Rugg, 2019). By this argument, the left inferior temporal gyrus is a region that is especially sensitive to cumulative lifetime experience with a perceptual category such as faces. Future research will be needed to test the validity of this proposal.

Lastly, we observed an own-age bias for the first presentation of unfamiliar faces in the right anterior hippocampus. Both young and older adults showed elevated BOLD signal for own-age relative to other-age faces. We stress that this finding of an own-age bias in the hippocampus requires replication within an experimental task that allows for a behavioral assay of own-age bias, such as recognition memory. Nonetheless, given the well-established role of the hippocampus in memory encoding (Eichenbaum et al., 2007; Kim, 2011), we conjecture that the present finding is relevant to the own-age bias that has been reported for recognition memory performance (Rhodes and Anastasi, 2012). There are several possible accounts of this finding, which are not mutually exclusive. One possibility is that it is easier to bind previously acquired personal or semantic knowledge about familiar individuals to unfamiliar own-age than to other-age faces. Alternately, own-age faces may more readily attract attention to discriminating facial features than other-age faces, in turn modulating encoding-related hippocampal activity (Aly and Turk-Browne, 2015, 2017; Uncapher and Rugg, 2009). Another possibility is that the present hippocampal effect reflects the formation of face representations with higher fidelity than representations of other-age faces (for reviews, see Ekstrom and Yonelinas, 2020; Yonelinas, 2013). Future studies that link the own-age bias in recognition memory to hippocampal effects promise to shed light on these and other possibilities.

There are limitations to this study. First, we did not replicate prior findings of own-age biases in regions such as the prefrontal cortex, insula, amygdala, and fusiform gyrus (Ebner et al., 2013, 2011a; Golarai et al., 2017; Wright et al., 2008; Ziaei et al., 2019). The reasons for this replication failure are unclear. One possibility is that task demands (e.g., passive viewing versus emotion recognition) contributed to the differences between the present and prior findings. Second, the own-age bias observed in the hippocampus did not have a behavioral correlate. While this is a potential limitation, we note that prior studies have reported own-age biases in neural data in the absence of analogous findings in behavioral data (e.g., Ebner et al., 2013; Ziaei et al., 2019).

In conclusion, we did not find evidence for own-age biases in face repetition suppression effects, which is arguably inconsistent with perceptual expertise accounts of the own-age bias. Of importance, a cluster in the right anterior hippocampus showed an own-age bias to the first presentation of unfamiliar faces in both young and older adults, which we speculate to be related to own-age biases in memory.

## Acknowledgements

This work was supported by an award to M.D.R. from the National Institute on Aging [grant number AG039103], and by awards to J.D.K. from the National Institute on Aging [grant number AGO49583] and the Aging Mind Foundation.

## Data Availability Statement

The behavioral data are available on the Open Science Framework (https://osf.io/bpk2d/). Un-threshold statistical maps from the voxel-wise fMRI contrasts are available on NeuroVault (https://neurovault.org/collections/11155/).

## Author Contributions

J.D.K. and M.D.R. designed the research, J.D.K. and N.H. performed data collection and analysis, and J.D.K., N.H., and M.D.R. wrote the paper.

## Abbreviation List

AC-PC: Anterior/Posterior Commissure
ANOVA: Analysis of Variance
BF: Bayes Factor
BOLD: Blood Oxygenation-Level Dependent
CVLT: California Verbal Learning Test
EPI: Echo-planar Imaging
fMRI: Functional Magnetic Resonance Imaging
FWE: Family-Wise Error
GLM: General Linear Model
MMSE: Mini-Mental State Examination
MRI: Magnetic Resonance Imaging
MPRAGE: Magnetization Prepared Rapid Acquisition Gradient Echo
SDMT: Symbol-Digit Modalities Test
WTAR: Wechsler Test of Adult Reading

